# Metappuccino: Large Language Model-driven Reconstruction of Sequence Read Archive Metadata for Cancer Research

**DOI:** 10.1101/2025.10.31.685769

**Authors:** Fiona Hak, Camille Marchet, Daniel Gautheret, Mélina Gallopin

## Abstract

**Motivation:** High-throughput RNA-sequencing has significantly advanced transcriptomic profiling in oncology. Millions of RNA-seq datasets have accumulated in public databases such as the Sequence Read Archive-SRA. However, fragmented, ambiguous or missing metadata can severely limit accurate cohort selection, introduce bias and delay discoveries.

**Results:** To address these issues, we introduce Metappuccino : a metadata enrichment tool based on a fine-tuned Mistral-7B-Instruct large language model with low-rank-adaptation (LoRA). Metappuccino can extract or infer 19 key metadata classes (e.g. organ, disease, cell type) from unstructured text. Fine-tuning was conducted with careful partitioning and training design to preserve the model’s generalisation capacity, reduce data leakage, and ensure robust, context-aware inference rather than memorisation. When possible, the inferred outputs are mapped to standardised ontologies, such as Cellosaurus, Disease Ontology and Uberon, to produce consistent metadata. As a result, the fine-tuned model achieves significantly improved class prediction accuracy over the base model, performing at least as well as recent large open-source models. Furthermore, it reduces inference time by up to at least two compared to the baseline models. As a pipeline, Metappuccino complements the LLM with well-established Natural Language Processing techniques from the literature to further improve performance. By enriching the metadata of under-annotated sequences, Metappuccino creates greater value from public RNA-seq datasets, with potential applications extending beyond oncology transcriptomics.

**Availability and Implementation:** The source code of Metappuccino is available on GitHub : github. com/chumphati/Metappuccino. The fine-tuned LLM, MetappuccinoLLModel, is available on Hugging Face : huggingface.co/chumphati/MetappuccinoLLModel. Both repositories are released under Apache-2.0 license.

## Introduction

High-throughput RNA sequencing (RNA-seq) has drastically improved our ability to characterize transcriptomic landscapes in various biological contexts, including oncology. At the same time, public repositories such as Sequence Read Archive (SRA) have accumulated massive data sets comprising millions of RNA-seq experiments, including more than 800 000 human RNA-seq samples produced with sufficient depth and quality for full transcriptome analysis between 2012 and 2025 (NCBI SRA Portal) (1). These data are highly valuable for the discovery of new clinical biomarkers and therapeutic targets, and for inferring transcriptomic signatures, i.e., patterns of gene or transcript expression that shed light on oncogenic mechanisms.

In spite of their great potential value, these databases remain largely underexploited. A major bottleneck is the lack of high-quality, available sample-associated metadata. Several issues have been identified while analyzing this metadata. (I) Sample metadata are retrievable through APIs (such as NCBI) with multiple fields that can be selectively queried. Still, some fields are absent from the API because they were not submitted, despite being present in the associated publication or stored in custom, non-standard fields. (II) Many fields are very often filled with useless values such as “unknown/missing/etc.” or are left empty. We found that 71.6% of the 192 fields across all run accessions in the NCBI API are left empty (Supplementary Figure 9).

(III) The fields can also be filled incorrectly : for example, age can be filled as a cell type. (IV) Finally, even when annotations are correct, they may be ambiguous because there are no normalized rules to submit some fields such as cell type or cell line. This can be very problematic for downstream analysis when, for example, trying to assemble a cohort. This can bias the extraction to always get the same well-annotated sample without exploiting the full richness of the databases (2; 3; 4). In this context, working with the current SRA metadata is very challenging and requires a substantial amount of human effort.

Several computational tools and methods have been developed to address these issues. Access and aggregation tools such as SRAdb and Grabseqs make SRA data easier to query and download but leave metadata untouched and inconsistent (5; 6). MetaSRA normalizes BioSample attributes by mapping them to ontologies with rule-based methods. It still remains sensitive to heterogeneity and ambiguities in metadata, which still prevents fully exploiting SRA’s potential (7). Newer systems move beyond field matching : SRA Database Navigator couples Natural Language Processing (NLP, automatic analysis of free text descriptions), Named Entity Recognition (NER, identifying and standardizing entities), and Network Analysis (graph-based linking and propagation) to reconstruct and enrich metadata for semantic clustering (grouping samples by meaning rather than keywords) (8). Txt2onto 2.0 uses large language model embeddings at prediction time to match unknown wording to known training terms, then applies Term Frequency–Inverse Document Frequency features (TF-IDF, weights a term higher when it is frequent in a document but rare across the document collection, highlighting discriminative words) to assign standardized labels (tissue, disease, etc). It is more robust to new phrasing but still bag-of-words, ignoring context and limited to training labels. (9). PredictMEE uses NER on unstructured BioSample text (e.g., titles, sample names) to infer missing attributes across 11 classes, achieving high accuracy, for example, Genus/Species (94%) and Condition/Disease (95%) (10). SGMC targets the geographic information by combining cloud queries, web scraping, and ChatGPT to standardize sequencing-institution and country fields across 2.3M SRA accessions, reaching 94.8% concordance for institutions and 93.1% for countries versus manual curation (11). ChIP-GPT fine-tunes a LLaMA model to answer ChIP-seq–specific questions, handling noisy fields and achieving 90–94% accuracy on targets/cell lines despite a narrow training scope (12). Most recently, Ikeda *et al*. (2025) showed that LLM-assisted extraction plus ontology mapping can outperform rule-based pipelines. From BioSample text, the method identifies cell line names and maps them to the cell line reference database Cellosaurus (13), surpassing MetaSRA both in accuracy and coverage (14). However, in all those methods, common gaps remain : (I) the use of small and not diverse datasets for LLM training that could lead to performance overestimation, (II) the reliance on constrained ontologies or rules/heuristics that limits the full exploitation of SRA’s potential, (III) the sensitivity to noisy text/lack of context understanding, (IV) the presence of manual or semi-automated steps, and (V) the limits in the quantity of extracted information with often maximum 1-3 classes (e.g. cell line). Most approaches also overprivilege the metadata stored in BioSample fields while underusing the study/BioProject context. This leaves out a lot of important information that is not stored in BioSample but that is still available elsewhere in SRA.

Building on those previous efforts, Metappuccino is the first hybrid and modular pipeline built around a fine-tuned Mistral model that goes beyond BioSample. It reads the full SRA record space by combining API fields, study and BioProject text, article abstracts, and optional user notes. The model both extracts values and infers missing ones, understanding medical context rather than relying only on rules. To limit typical LLM issues such as hallucinations, it pairs the model with classic rule-based NLP similar to MetaSRA, then uses biomedical ontologies to harmonize terms and cut ambiguity. Unlike tools limited to a few attributes, Metappuccino now extracts 19 classes, including standard SRA fields and additional ones, manually defined, useful for downstream cancer transcriptomics (Figure 1). These classes are not fixed : the training setup and pipeline design allow new ones to be added on request. Metappuccino can be applied to any kind of data beyond transcriptomics in oncology, as long as outputs from irrelevant classes are ignored.

**Figure 1.**
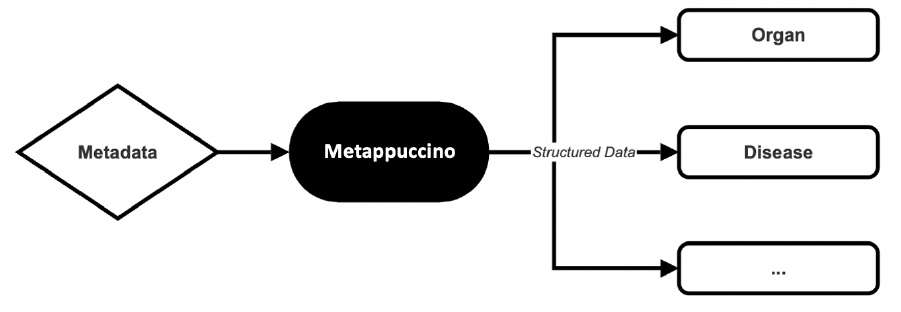
From Raw Metadata to Structured Information.

## Methods

### Metappuccino pipeline

The Metapuccino pipeline v.1.0.0 can extract 19 different types of information (discrete or open-vocabulary) from a context, which we will refer to as classes (Table 1).

**Table 1.** Metappuccino extracted metadata fields.

|  |  |
| --- | --- |
| Study accession number : | unique study ID. |
| Instrument Platform : | sequencing technology family (e.g. Illumina). |
| Number of base pairs : | total number of sequenced bases. |
| Library strategy : | assay type (e.g. RNA-seq). |
| Library selection : | library enrichment strategy. *Discrete values* : polyA, inverse rRNA, hybrid selection, small RNA, other. |
| Sequencing source : | sampling granularity of RNA-seq. *Discrete values* : bulk, single cell, spatial. |
| Biopsy site : | organ, tissue, body part, or fluid from which the sample was extracted. |
| Biopsy type : | clinical context of the sample *if cancer related*. *Discrete values* : primary, metastasis, blood. |
| Cell line : | standardized name of cultured/immortalized line. |
| Cell type : | cell identity, that share morphological or phenotypical features (e.g. neuron, fibroblast, CD8 T cell, melanocyte). |
| Organ : | organ affected (distinct from the sampling site in biopsy site). |
| Disease : | specific associated condition (or healthy). |
| Treatment : | intervention applied (drug, procedure, implant, event, etc.), including planned pre-treatments. |
| Treatment time : | timing relative to the applied intervention (e.g. pre, on, post, or a duration). |
| Response : | clinical or experimental outcome. *Discrete values* : no treatment, unknown, stable, progressive, success. |
| Age : | donor age if human. |
| Sex : | donor sex if human. |
| Ethnicity : | donor ethnicity if human. |
| Is cancer : | whether the disease is cancer-related. *Discrete values* : True, False. |

#### 1. SRA Extraction

Starting from a list of SRA-run accession numbers, Metappucino retrieves for each run an SRA XML file via the NCBI E-utilities API (2. XML, 3). This run-level record is structured into four blocks (<STUDY>, <SAMPLE>, <EXPERIMENT>,<RUN>) describing, respectively, the project or study, the sample,the experimental setup, and the sequencing run. The XML file is parsed into (I) reading study metadata from <STUDY> (title, first public date, project identifiers, abstract, etc.) (Figure 2. Study information), (II) reading sample metadata from <SAMPLE> (title, free-text description, age, biological source, etc.) (Figure 2. Special fields), and (III) getting a BioSample accession (SAMN…) from cross-references XREF_LINK and EXTERNAL_ID (if absent, a regular-expression search within the SRA XML is attempted). Using the resolved accession, the corresponding BioSample XML file is downloaded (Figure 2. Biosample attributes). In parallel, an SRA API call returns values for fields that may be relevant to the targeted classes (details in *Supplementary. Extracted fields from the NCBI API during pipeline download*). All retrieved items (SRA XML, BioSample XML, and SRA API fields) are finally merged into a tab-separated file with one row per run. Metappuccino also accepts an optional tabular file with a required run accession column and any number of user-supplied text-metadata columns, which is useful if a user has access to additional data to complement SRA information.

**Figure 2.**
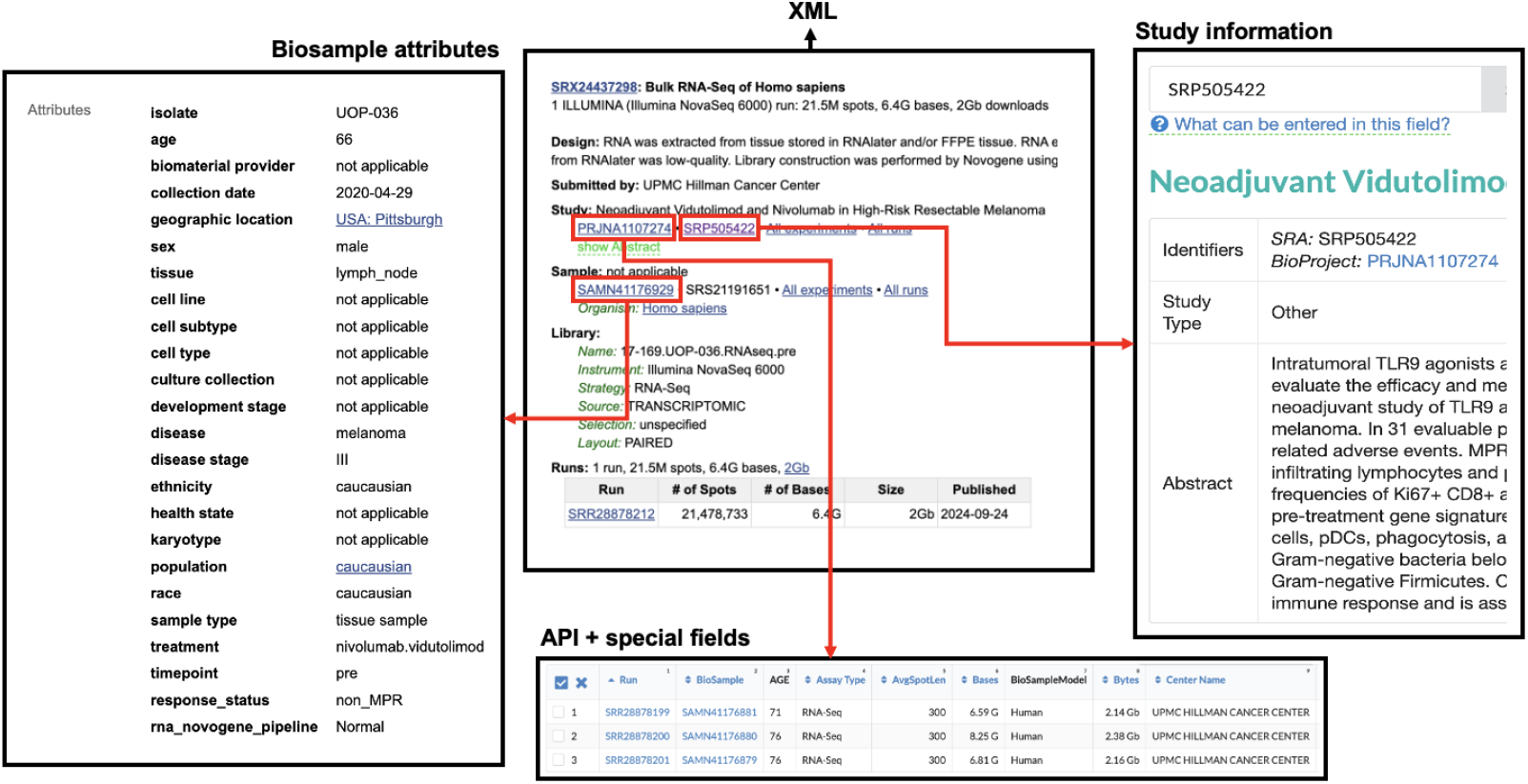
Example of SRA inputs collected by Metappuccino for SRR28878212. BioSample contains sample-centric information filled by the submitting team. Study information provides project-level metadata shared by all samples. The SRA XML aggregates and links study, sample, experiment, and run information, and it may include author free-text fields not exposed in some views/APIs. Some information appears in multiple sources, while other is specific to a single source.

#### 2. Pre-processing

The preprocessing stage has two main steps (Figure 3) :

**Figure 3.**
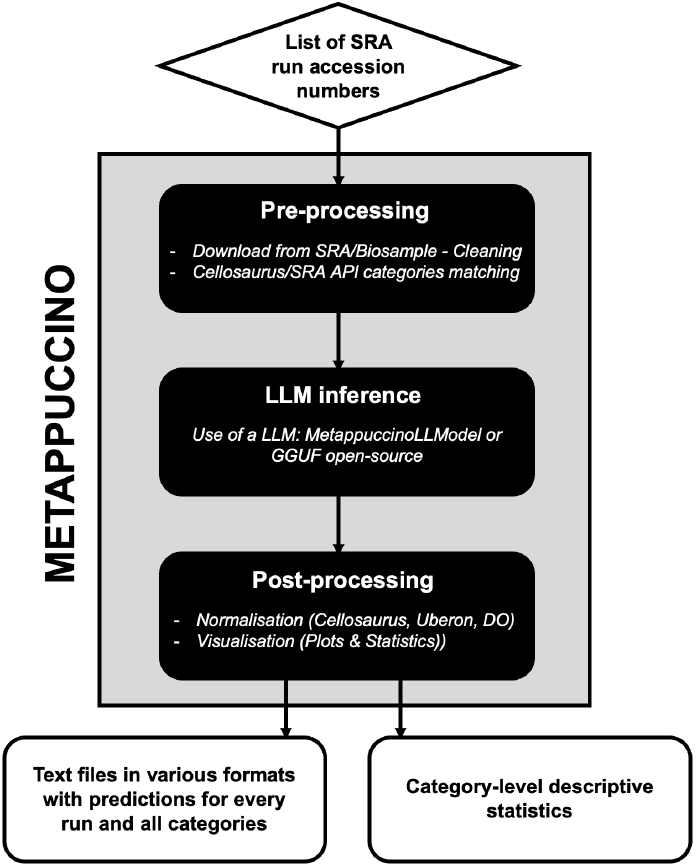
Metappuccino Workflow. Metappuccino first downloads and pre-processes SRA run records to extract as many class values as possible with rule-based NLP. LLM inference then completes any missing fields. Post-processing uses medical dictionaries and ontologies to infer related fields from new predictions and to normalize terminology. Outputs are tables (various formats) with per-run, per-class values, plus descriptive statistics.

—Turning the raw SRA and BioSample downloads into a clean, compact paragraph-style summary for each run. First, a cleaning part converts the XML files into paragraphs. Then, the text is summarized only if needed (over 2000 tokens). In that case, obvious identifiers and boilerplate are removed, duplicates are deleted, and rare informative phrases and words are kept. This part is important in order to avoid giving the LLM too long contexts, which reduces performance and increases hallucinations, but the variety of the biological context has to be maintained.

—Extracting values for the 19 classes from the raw downloads whenever this is supported by our rule-based NLP methods : (I) When a class name appears as an explicit XML tag or a BioSample attribute, the value is directly extracted, including common aliases. (II) When a class exists only in free text, it is extracted by phrase matching using a dictionary build from Cellosaurus (13), where controlled names and synonyms are lowercased, purged of generic terms, and compiled into a spaCy *PhraseMatcher* data structure (the text is parsed once, exact spans are returned, candidates are deduplicated while preserving order, and a lightweight regex fallback catches missed alphanumeric variants). (III) When data is missing for `library_selection, sequencing_source and biopsy_type,` simple heuristics try to infer them when their values are explicitly stated or via a list of their synonyms. (IV) When a cell line is found by one of the previous methods, the Cellosaurus (13) database is used, first to normalize the extracted term if it is a synonym to a canonical cell line name, and then to fill the following classes based on its knowledge of the cell line : `biopsy_site, cell_line, cell_type, organ, disease, age, sex, and ethnicity.` Finally, in addition to Cellosaurus, a normalization step maps diseases to Disease Ontology (15) names and identifiers (diseases database) and organs or biopsy sites to Uberon (16) names and identifiers (species anatomy database). Values obtained by steps (I)–(III) are validated before acceptance as follows : (I) *type and range* checks (e.g. age must be numeric within plausible bounds, sex in {male,female,…}, etc.); (II) *ontology* checks (disease, organ, cell_type, and biopsy_site must map to valid terms in Disease Ontology, Cellosaurus, or Uberon); *cross-source consistency* across SRA XML, BioSample attributes, and API tables, with agreements preferred and disagreements flagged for downstream LLM review. Any value failing a check is rejected from the curated table. All extracted and normalized values are merged in a temporary file with one column per class and one run per row. Those values will not be overridden by the following LLM completion as they are considered true.

#### 3. LLM Inference

LLM inference uses the prepared summary and the ambiguity notes from preprocessing to predict 15 classes out of 19. This is because four classes (Study accession number, Instrument Platform, Number of base pairs and Library strategy) are always populated in their corresponding NCBI API fields. Two interchangeable LLM backends are supported : (I) **Hugging Face Transformers + PEFT (LoRA)**, our MetappuccinoLLModel mode, where Transformers is the PyTorch stack and PEFT (Parameter-Efficient Fine-Tuning) updates only small add-on modules instead of all weights (17). LoRA (Low-Rank Adaptation) inserts trainable low-rank matrices into attention/MLP projections (17). We provide one LoRA adapter per class (see *Data Availability*) and activate the relevant adapter at inference on the base model `mistralai/Mistral-7B-Instruct-v0.3.` (II) llama.cpp (github.com/ggml-org/llama.cpp), which runs local GGUF models (a quantized, CPU/GPU-friendly format downloadable from Hugging Face) without PEFT, letting the user swap in any compatible model, including non open-source ones. This mode exists to give users the final say on the model, allowing easy model switching and enabling fully local deployment.

For class extraction, the prompt was developed empirically : different wording and tags were tested, the outputs were checked, and the structure that gave clear, direct answers was kept (Table 2). For each run and class, the run accession *(a)*, the summary and the ambiguity notes *(b)*, the class definition *(c)*, other information *(d)* are included into the prompt template. The system outputs one JSON file per run containing all predicted class values and two confidence scores (Supplementary Figure 17) : the mean negative log-likelihood (NLL) and the perplexity (PPL) which quantify the uncertainty of a model when predicting the next token in a sequence. For each class, the model outputs a short sequence of *T* tokens, written *y*_1:*T*_. At each step *t*, it assigns a probability *p*_*t*_ to the token it emits. The *token-level uncertainty* is computed by *s*_*t*_ = *−* log *p*_*t*_. Then, the average of *s*_*t*_ is taken : 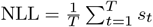 Perplexity is defined as 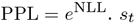 and kept to facilitate interpretability. Lower NLL and PPL indicates higher confidence.

**Table 2.**
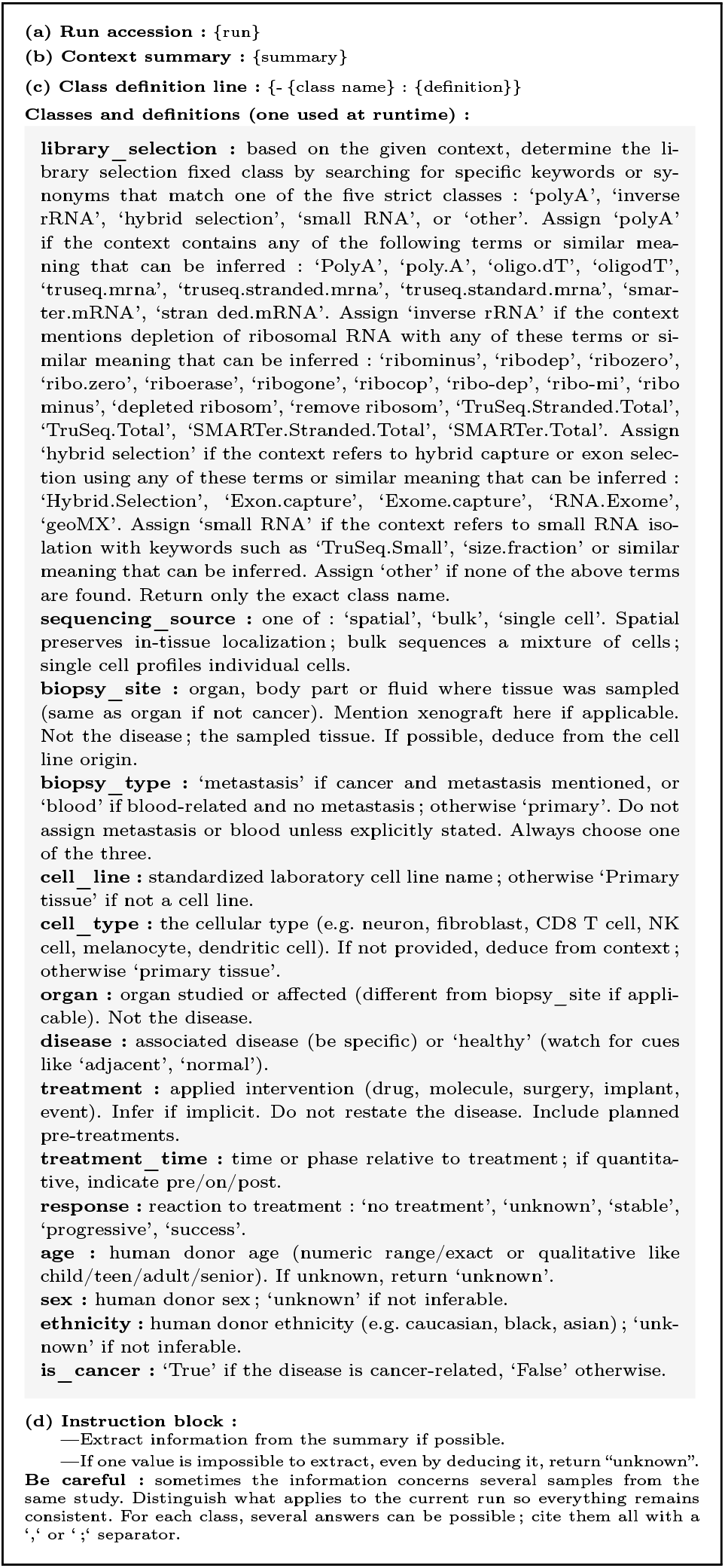
Dynamic prompt template used for LLM inference. (a) run accession, (b) context summary, (c) the class definition line (one picked at runtime), and (d) the instruction block are injected for each run and class for individual inference.

#### 4. Post-processing

The post-processing stage converts the curated metadata extracted at preprocessing into a single table per run, using rule-based NLP and the LLM predictions, normalized with Cellosaurus, Disease Ontology and Uberon. Values extracted at preprocessing are locked, and only missing fields are complemented with the LLM inferences. Additional cell lines detected during LLM inference are normalized with Cellosaurus and can propagate consistent attributes (e.g. disease, cell type, biopsy site). Free-text organ/disease/biopsy terms are normalized to preferred names and mapped to stable codes, with light cleaning (synonyms, formatting). The final dataset is exported into multiple formats (CSV, TSV, XLSX, Parquet, JSON, Feather), alongside confidence tables, colored HTML views, and summary graphics to support quick quality checks and downstream analysis.

### MetappuccinoLLModel

A task-specialized fine-tuned model is provided with Metappuccino : MetappuccinoLLModel, trained to outperform available open-source LLMs on our task.

#### Fine-tuning

`Mistral-7B-Instruct-v0.3` is our backbone, selected for its fastest native inference among the open-source models we tested (see *Baseline-LLM selection*). We freeze the base weights and train lightweight LoRA adapters (17), one per class, to keep deployment fast, modular, and memory-efficient. Training uses the same prompt format as inference, runs at 16-bit mixed precision on two A6000 GPUs (80 GB VRAM), and is designed to teach the model where to extract each classes’ information within different context structures,maximizing use of context while reducing hallucinations and extraction errors, while producing concise outputs. All hyperparameters and settings are documented in the Hugging Face configs of the released adapters (https://huggingface.co/chumphati/MetappuccinoLLModel). Further details of the training method are available in the *Supplementary Material, MetappuccinoLL-Model* : *In-Depth Methods, Training procedure and final adapters selection*.

#### Data for Training and Evaluation

Because well-annotated SRA data are limited, especially for our target classes, we created large scale synthetic texts based on Metappuccino summaries, that closely matched real inputs. We found that the best results per class used about 600 to 3000 training samples and about 200 to 500 validation samples, with balanced classes. For open-vocabulary classes, values are strictly different across training, validation, and test sets to promote generalization, not memorization. Models are evaluated with three test sets per class : **ID (identical) with 400 samples similar to training data, OOD (out of distribution) with 400 samples very different from training data**, and **MID (middle) with 1200 samples that mix both**. Full details on data generation and splitting are given in *Supplementary Material, MetappuccinoLL-Model* : *In-Depth Methods, Data Generation*. To complement synthetic data, we randomly selected 400 human RNA-seq SRA records and 100 melanoma records, also randomly selected among melanoma annotated samples, that we manually annotated for all target classes. These annotations are available in the Data section of the GitHub repository. The 100 melanoma records were annotated twice, once using only SRA information to fairly evaluate the LLM given the same context it sees, and once with added Cellosaurus knowledge for the *Study case with Metappuccino, 100 melanoma samples*, which shows how pre and postprocessing can improve performance alongside LLM inference.

#### Data Leakage

Contexts and values were disjoint across train, validation, and test splits, which reduced direct overlap. To further verify the absence of leakage due to overly similar contexts, two embedding-based diagnostics were computed per class and per split (Figure 4) : (I) A distributional shift was measured with the squared Maximum Mean Discrepancy (MMD). This measures how far apart the overall shape of a dataset’s points is from the training points : smaller values mean the two distributions are more similar. The comparison uses distances based on cosine similarity, and a Gaussian smoothing factor whose width is automatically chosen using the median of all pairwise distances. (II) The near-duplicates : for each dataset item, the most similar training item is found using cosine similarity. If this maximum similarity is very high (more than 0.95), the test item is counted as a near-duplicate (ND). ID splits show low MMD as expected (close from training by nature), MID are intermediate, and OOD and real-world tests exhibit larger MMD, consistent with intentional distributional shift and embedding distributions (Supplementary Figure 14). Depending on the class, MID and ID can show very similar MMD, and MID can even be lower. This is expected because MID usually has almost 3 times more samples, which reduces the variance of the empirical MMD estimate, and in some classes MID is genuinely closer to the train distribution. Near-duplicate rates are essentially zero in test splits, with only small residuals on a few validation sets (e.g. age), which aligns with constrained phrasing in some classes. Those observations reflect heterogeneous training distributions, especially when real SRA contexts are part of the training sets, which expands the coverage of the training embedding space and can raise baseline MMD without implying leakage. Overall, the diagnostics support minimal data leakage and a clean separation between in-distribution and shifted test conditions.

**Figure 4.**
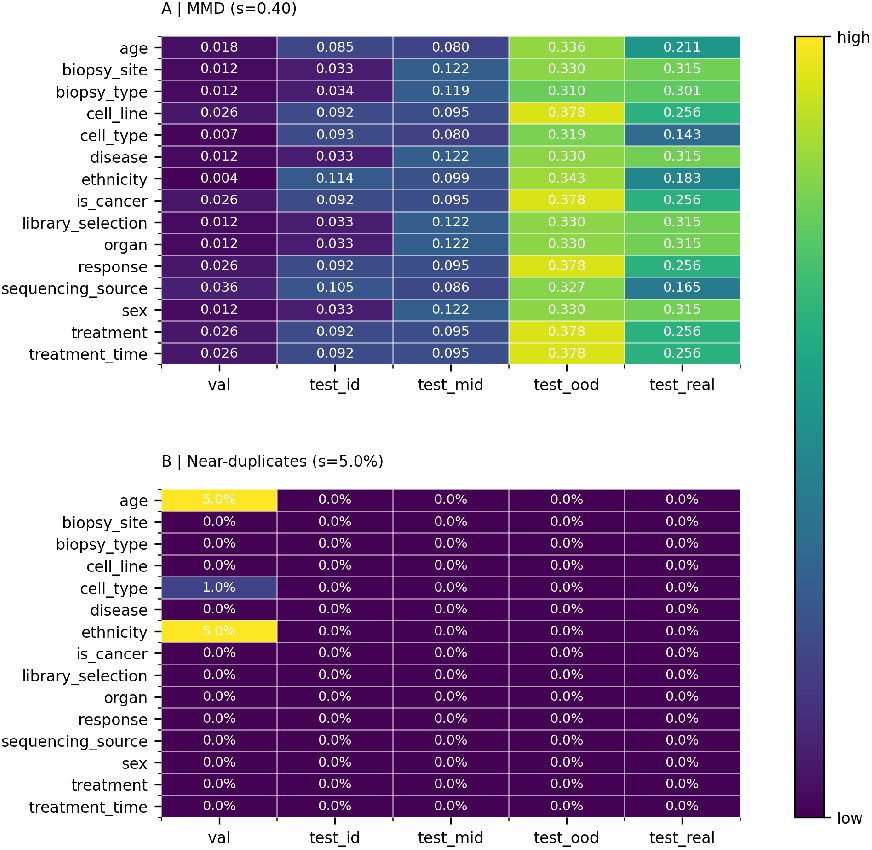
Data leakage evaluation between train, validation, and test sets. *MMD (A)*. Small values mean that the set has similar distributions to the train. *Near-duplicates (B)*. The table reports the percentage ND, so values near zero indicate that copy-like items are absent.

### Benchmarking

The performance of Metappuccino and MetappuccinoLLModel was assessed against commonly used open-source LLMs.

#### Baseline-LLM selection

The following instruction-tuned open-weight models were chosen as reference baselines for this study : Mistral-7B-Instruct-v0.3 (18; 19), Llama-3.1-8B-Instruct (20), Llama-3.1-70B-Instruct (20), DeepSeek-V2-Instruct (21; 22), Gemma-3n-E4B-Instruct (23), and Qwen3-30B-A3B-Instruct (24). These models span a spectrum from small, efficient LLM variants suited to modest hardware (Mistral-7B, Llama-3.1-8B, Gemma-3n-E4B) to larger, higher-capacity systems for higher accuracy and longer-context reasoning (Llama-3.1-70B, Qwen3-30B, DeepSeek-V2). ‘XB’ designates the number of parameters of the model. All are instruction-tuned to follow task prompts reliably. They were all downloaded via Hugging Face on their official repositories. They were selected to evaluate the most widely used open-source originals. The latest versions were tested, and only those that produced responses consistent with our prompt structure were retained. Q8 quantization was used for Llama-3.1-70B to meet memory and throughput constraints (2 GPU = 48GB x2 VRAM, NVIDIA-A6000). All other models were loaded unquantized in floating-point : FP16 by default. The same prompt and datasets were used for inference with all the models to produce comparable results.

### Performance Evaluation

A hybrid evaluation was implemented to handle both classes requiring : (I) context-tolerant matches (e.g. organ : *oral cavity* vs. *mouth* = True) and (II) exact matches (e.g. is_cancer : true/false).

#### (I) Soft accuracy for context-tolerant classes

Given a set of *N* samples, 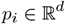is the embedding of the model’s prediction and 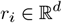the embedding of the reference metadata for sample *i*. These embeddings are obtained by encoding each text string with a sentence-transformer model (*pritamdeka/BioBERT-mnli-snli-scinli-scitail-mednli-stsb*), which maps semantically medical similar terms to nearby points in the *d*-dimensional space. The cosine similarity between *p*_*i*_ and *r*_*i*_ is

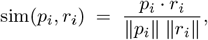

where the numerator is the dot product and the denominator the product of Euclidean norms. A higher value (up to 1) indicates stronger semantic alignment. If sim(*p*_*i*_, *r*_*i*_)*≥ τ*, the prediction is counted as correct. Thresholds *τ* were set empirically, searching where the right/wrong switch occured in the comparison reference/predictions (e.g. ref=‘True’, prediction=‘False’, score=0.3, *τ* = 0.2, answer=‘matching’. Here, the answer ‘matching’ is wrong, higher thresholds are needed to get answer=‘non matching’). Finally we founded that : *τ*_default_ = 0.40, *τ*_disease_ = 0.42, *τ*_organ_ = 0.37, *τ*_ethnicity_ = 0.30, *τ*_treatment_ = 0.33, *τ*_treatment_time_ = 0.35. The default applies to all other classes. The semantic accuracy at threshold *τ* is then

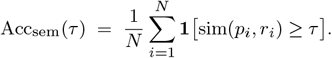

#### (II) Classical accuracy for exact-matches classes

For classes requiring strict identity, strings are lowercased and punctuation is removed before comparison, and a prediction is correct only if the normalized strings are identical.

### System requirements

Metappuccino runs with PBS and Slurm schedulers, and on local Linux/MacOS systems. With MetappuccinoLLModel, we recommend 8 CPU cores and one GPU (at least 40 GB VRAM); CPU-only execution is supported but significantly slower (see the project README, release v1.0.0, commit 136, for technical details).

## Results

### Predictive performance of MetappuccinoLLModel vs standard open-source LLMs

The performances of DeepSeek, Gemma, two Llama LLMs of different numbers of parameters, Qwen3 and untuned Mistral were compared to MetappuccinoLLModel. Two trivial classifiers were also included : the “Majority class” that reports the accuracy obtained by always predicting the most frequent label in a class; and the “Unknown class” that reports the accuracy obtained by always answering unknown. This way, majority-class bias and a tendency to answer “unknown” frequently by a model can be detected more easily.

### Accuracy achieved on the synthetic sets

On ID sets, MetapuccinoModel outperforms all untrained models for all classes with an accuracy > 80% for 8 of them, and never below 60% (Figure 6.A.). These results are expected because the ID test dataset is similar to the training dataset (Supplementary Figure 14).

**Figure 6.**
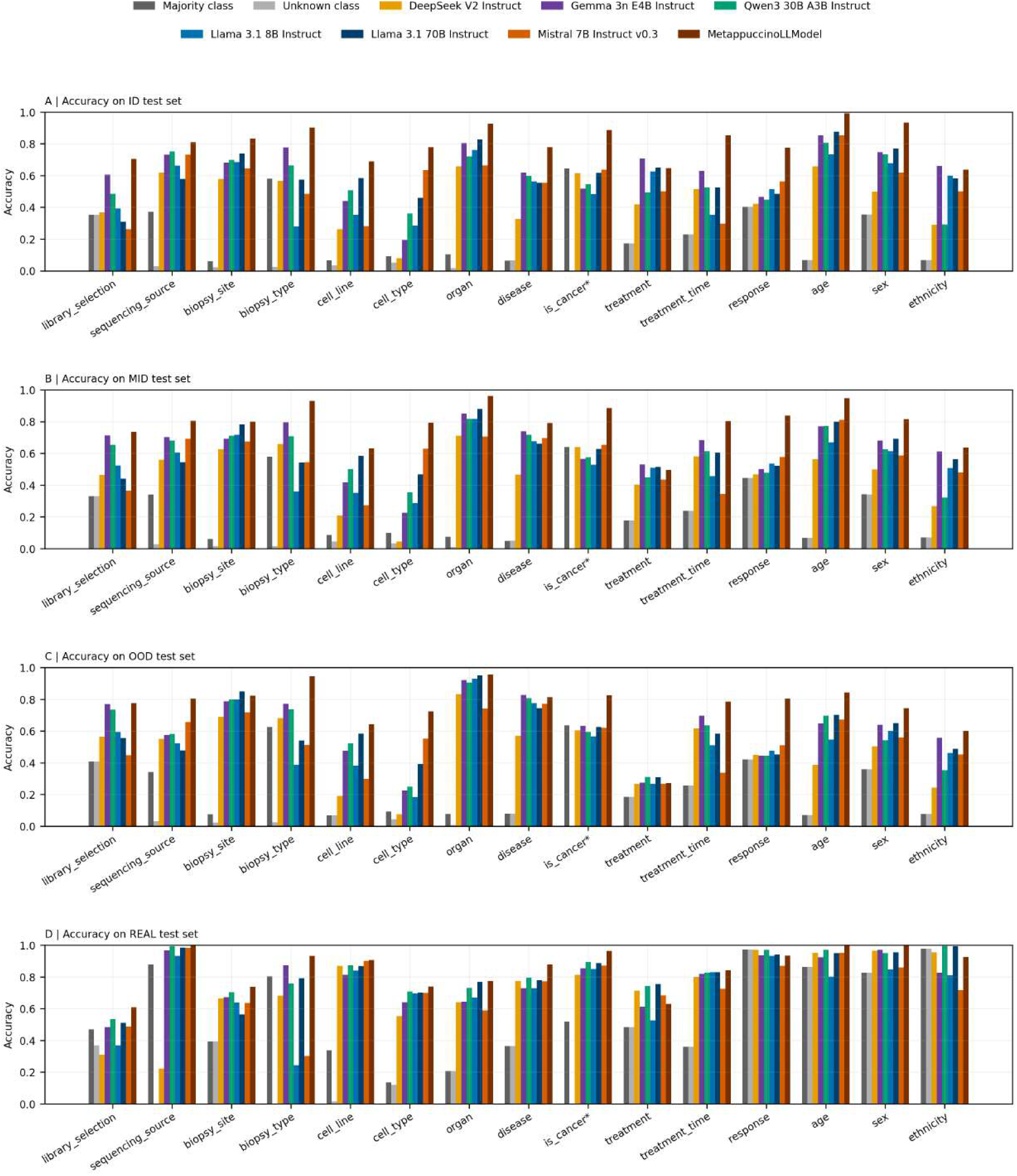
Per-class accuracy across the different test splits. A–C. Accuracy on synthetic datasets constructed from an initial set of 2000 samples, split by embedding proximity to the training set : ID (400 samples) for close points, OOD (400 samples) for distant points, and MID (1200 samples) for intermediate points. D. Dataset of 500 samples randomly extracted from human RNA-seq SRA sample metadata, excluding any samples used for model training.

To assess generalization under distribution shift, we evaluated models on the MID split, which contains contexts slightly structurally different from the train set (length, vocabulary, etc.) (Supplementary Figure 14). Accuracy drops moderately, yet remains high and above both untuned Mistral and the strongest baselines for almost all classes (except treatment) : 11 classes have around 80% accuracy or greater (Figure 6.B.). Model rankings mirror those on ID, indicating predictable generalization; MetappuccinoLLModel remains first in classes where context understanding is important (e.g. biopsy type, treatment time, response). Because MID has more samples than ID (1200 vs 400), slight gains occur — for instance, organ at 0.9617 (MID) vs 0.9275 (ID) — partly due to reduced variance and the split procedure from a single balanced pool by embedding proximity (Supplementary Figure 12), which can mildly overrepresent easier cases. Overall, performance degrades slightly in a predictable way with distance to the training set or stays stable depending on the class.

One important aspect to verify is that MetappuccinoLLModel does not rely on memorization and can process inputs very different from its training data. In the OOD split, the strongest shift, accuracy remains robust (Figure 6.C.). Nine classes still reach > 80% accuracy. MetappuccinoLLModel still outperforms all models except for biopsy site (Llama-70B) and disease (Gemma). Treatment remains poorly predicted at 0.2725 across all models, indicating an intrinsically hard class. Large margins over majority and unknown classifiers are preserved. For example, treatment time 0.785 vs 0.2575 for the majority/unknown. Therefore the model is not biased toward those default answers. Because open-vocabulary values are disjoint across datasets (e.g. if “lung” is in the class “organ” for the training set, it cannot be in any test set), these results cannot be attributed to memorization.

### On real-context sets

To verify performance on unmodified real data, we evaluated 500 manually annotated SRA records across all 15 classes. It can be observed that when metadata is explicit in the given context, strong models all perform equally well. For instance classes age and cell line are often stated explicitly (e.g., age = x years). When information is this clear, any LLM can extract it reliably (Figure 6.D.). Where wording varies and domain knowledge matters (library selection, biopsy site, biopsy type, organ, disease, cancer flag, and cell type), MetappuccinoLLModel stays ahead. Treatment remains poorly predicted and shows no clear fine-tuning gains, especially when context differs from training. A key limitation of this real dataset is class imbalance : many classes have a dominant answer, such as sequencing source (often bulk), and response with 97.4% of cases annotated as unknown, which mechanically inflates baseline accuracies because untuned LLMs often default to “unknown” or prompt defaults when the information is not clearly stated (Figure 6.D.). Along with natural variation in how easily values can be found, this explains why gaps between MetappuccinoLL-Model and untuned models are smaller than on synthetic data. Nonetheless, MetappuccinoLLModel maintains strong predictive performance across all classes on real data.

### Main prediction error types

The prediction errors made by each model for each classe were investigated based on the errors made on all the test sets (Figure 6).

#### MetappuccinoLLModel

About 30.6% of errors are abstentions (“unknown”). The remaining 69.4% are substitutions, as follows : library_selection = other (8.9%), cell_line = primary tissue (3.9%), biopsy_type = primary (3.1%), is_cancer = true/false (4.3% total), sequencing_source = spatial/bulk/single-cell (5.5% total), treatment_time = not applicable (2.0%), and response = stable/progressive (2.3%); the rest are many low-frequency substitutions.

#### Mistral

About 42.3% of the errors are abstentions. Among substitutions, the dominant patterns are library_selection = polya (8.6%), cell_line = primary tissue (6.1%), is_cancer = true (2.7%), sequencing_source = bulk (2.4%), with smaller but recurring cases like sex = male (0.8%), sequencing_source = spatial (0.6%), biopsy_type = primary (0.6%), and cell_type = primary tissue (0.5%).

#### Other baseline LLMs (average)

About 58.6% of errors are abstentions. Substitution patterns are more diffuse; the largest include cell_type = primary tissue (4.5%), is_cancer = true (2.7%), library_selection = polya (2.1%), sequencing_source = bulk (2.1%), and biopsy_type = primary (2.1%), with smaller contributions from library_selection = other (1.0%), sex = male (0.9%), is_cancer = false (0.7%), and sequencing_source = spatial (0.7%).

In conclusion, thanks to fine-tuning, MetappuccinoLLModel abstains less, even if it keeps some error patterns from its Mistral backbone. When Metappuccino is wrong, it is more likely to make a plausible, structured substitution than to fall back to abstention.

### Inference time across models

We measured inference times for 2000 samples on one NVIDIA A6000 GPU with a context size of minimum 589 tokens, maximum 3500 tokens and a mean of 2337 tokens. MetappuccinoLLModel is the fastest model, with a median of around 4 seconds per sample and a tight dispersion, with most runs processed between 3 and 8 seconds (mainly depending on context size). Untuned Mistral is slightly behind with a median of around 6 seconds. Llama-3.1-8B and Qwen have intermediate performance with medians of 13 and 15 seconds. DeepSeek has a median of around 14 seconds Llama-3.1-70B moves to a median of 23 seconds. Gemma is even slower, with a median of 26 seconds. Finally, MetappuccinoLLModel is clearly the fastest and most stable, with intermediate sizes offering compromise and very large models being more expensive and more variable in inference time (Figure 7). On CPU, the inference time for MetappuccinoLLModel averages 54 seconds, compared to over 15 minutes for Llama-3.1-70B.

**Figure 7.**
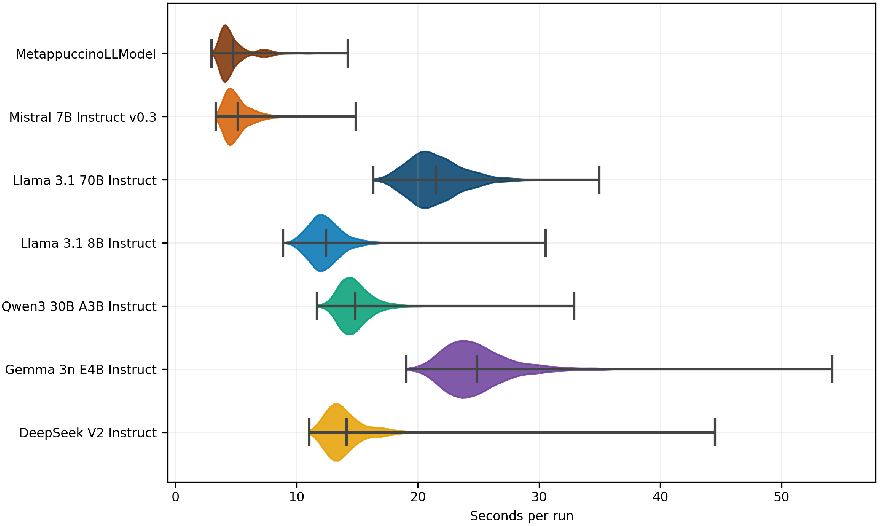
Distribution of per-sample inference times (seconds) for each model. The central tick marks the median, whiskers indicate the outer range.

### Contribution of MetappuccinoLLModel to Metappuccino : a case study with 100 manually annotated melanoma samples

Class values are exactly extracted when possible by Metappuccino’s rule-based preprocessing and post-processing (see *Methods*). We tested whether these deterministic methods alone could match or surpass LLM inference. In Figure 8, accuracy per class is computed only on runs whose reference is not “unknown” or “not applicable,” comparing “Preprocessing Only” to “Pre/post-processing + MetappuccinoLLModel” (full pipeline). LLM-only inference was not evaluated here, as it lacks ontology access, especially to Cellosaurus, and is therefore not comparable without the post-processing dictionary match. Results show consistent gains with the full pipeline. The largest LLM improvements appear for cell type, treatment, treatment time, organ, and disease. Library selection improves slightly, although LLM predictions there remain less reliable than for other classes. Overall, LLM processing improves every class, and the full pipeline delivers the strongest performance.

**Figure 8.**
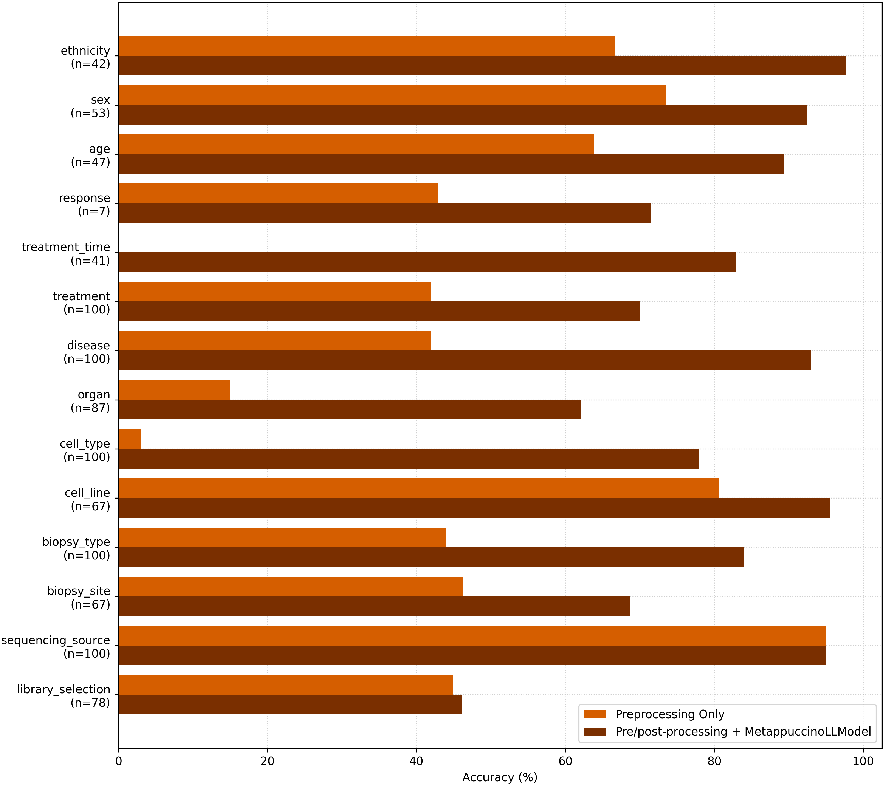
Metappuccino performances on the known values of the 100 melanoma-real dataset. Grouped accuracies (%) for “Preprocessing Only” and “Pre/post-processing + MetappuccinoLLModel”. Each bar is computed only over non-unknown references for that class (n).

## Discussion

### MetappuccinoLLModel training

We tested two training setups, one model learning all 15 classes with a single loss (negative log-likelihood of the true label under the model’s predicted probabilities, allows to measure the performance progression during training), and separate training for each class. The single loss setup was unstable, when a few classes improved, others stalled or got worse because the hardest ones held back the rest. At inference, asking for one class at a time worked better than asking for all 15 at once, no matter which LLM was used. We therefore use per class training : it reduced interference, helped each class to specialize. Overall, performance depended most on data quality, some classes improved with more varied phrasing in the training texts, others with cleaner, hand checked examples.

### Why are there performance differences between classes ?

Performance varies with the class, the clarity of the prompt, and the quality of the input. Discrete classes have few values, so the model can learn patterns and be guided to detect them rather than generate text. Open vocabulary classes rely on semantic matching to learn, so we must define what counts as correct. For example, “TB” versus “tuberculosis” can be considered as correct, but special attention is needed to prevent drift and hallucinations during training caused by allowing different values than the target. Data composition also matters. In SRA or BioSample,the target information to extract is rarely written clearly, or it follows writing patterns that differ across studies. For example, the class “sex” is almost always stated explicitly, while the class “organ” may be absent yet inferable from a disease (e.g. lung inferred when pneumonia is present). Difficulty increases even more with text diversity and distance from training examples, which explains gaps between ID, MID, and OOD tests : when learned patterns are missing, similar performance is harder to achieve. For example, if the model does not know all the ways to express “bulk”, it may only recognize patterns seen during training. Moreover, prompt engineering strongly affects results. Some classes are hard to annotate even for humans. For example, treatment time is often unclear (e.g. whether a time is pre, post or on treatment), or contexts may describe treatments applied during a study without confirming that they apply to the specific sample that has to be annotated. Such biases make extraction difficult overall, and LLMs do not outperform humans on such cases.

Regarding prediction errors, MetappuccinoLLModel often makes near-miss substitutions that a human might accept, for example, “surgery” instead of “lung surgery”, whereas other models more often produce unrelated substitutions, an effect that reflects progress with the fine-tuning improving the fallback to simple defaults answers.

### A standard LLM with adapters

We chose an LLM approach with per-class LoRA adapters because it is flexible : adding a new class does not require retraining the entire model but just creating a new dedicated adapter and train it locally without affecting the others. This reduces cost, limits regressions, simplifies maintenance and allows quick evolution of the tool. This solution also avoids using multiple models for each class during inference, which would be inefficient in practice : even if some baseline LLMs sometimes achieve a similar or slightly higher score on a given class, MetappuccinoLLModel keeps the advantage across all class, with a homogeneous and stable behavior.

### Already open-source fine-tuned LLMs

We tried to use already fine-tuned models on medical data but they are often optimized for other tasks (clinical QA, summarization), use training sets that are hard to audit, and can introduce bias. Moreover, on our task, they did not perform better than the general LLMs.

### Evaluation without bias

Prediction accuracy depends on the dataset : how much information is present, how varied the phrasing is, and how balanced the values are, so an absolute unbiased metric is hard to give. Still, the trend is clear, MetappuccinoLLModel generally outperforms baselines models, and model rankings stay stable as context patterns and values change, showing good generalization. A baseline model can match or beat it on a specific class, but the overall advantage remains. Pre and postprocessing further improve answers, and normalization with Cellosaurus, UBERON, and Disease Ontology keeps outputs consistent and medically grounded. The weaknesses of rule-based NLP are offset by the strengths of the LLM, and vice versa. Taken together, *Metappuccino* delivers the strongest metadata reconstruction to date.

### Limitations : treatment and library selection

Treatment and library selection remain harder to predict, they depend more on pre and postprocessing. Treatment details are often diluted in study text, the model may mix treatments across samples or list all study treatments instead of the sample’s. Library selection fields blend protocols, abbreviations, and technical notes, which confuses any LLM and even humans (e.g. when ribodepletion and polyA selection are described in succession). These limits come not only from the model, but also from unclear inputs and, at times, the prompt.

### Future directions

The use of adapters make Metappuccino easy to extend to new classes and domains. It is currently specialized for human cancer metadata, yet already usable beyond oncology when class-specific fields are ignored, and it can be expanded to bacteria, plants, or other organisms by adding new adapters. Enriching the inputs given to Metappucino, beyond SRA metadata, in order to reduce the number of unanswered predictions is another path for improvement. Users can already provide their own supplementary metadata documents, but exploring automatically new sources would give, we believe, a significant boost. We envision using longer documents such as full research articles, and even switching to other extraction methods beyond LLMs, such as Vision Transformers.

Pending these improvements, Metappuccino already offers a robust, scalable solution for extracting and completing metadata from a simple list of SRA run accessions, combining rule-based NLP with LLM inference, requiring a few seconds per run on GPU while remaining operable on minimal CPU infrastructure. While extracted classes are currently oncology-centered, Metappuccino is applicable across diverse data types.

## Author Contributions

Study design : FH, DG, MG, CM; manuscript writing : FH, DG, MG, CM; statistical overview : MG; software development : FH.

## Data Availability

Metappuccino and all the materials used in this study including the fine-tuning and evaluation scripts are available in our GitHub project : https://github.com/chumphati/Metappuccino. The associated fine-tuned adapters are available on Hugging Face : https://huggingface.co/chumphati/MetappuccinoLLModel.

