## Supplementary Material for "Metappuccino: Large Language Model-driven Reconstruction of Sequence Read Archive Metadata for Cancer Research"

### Supplementary Material
### 1. Initial SRA composition of fields extracted by Metappuccino

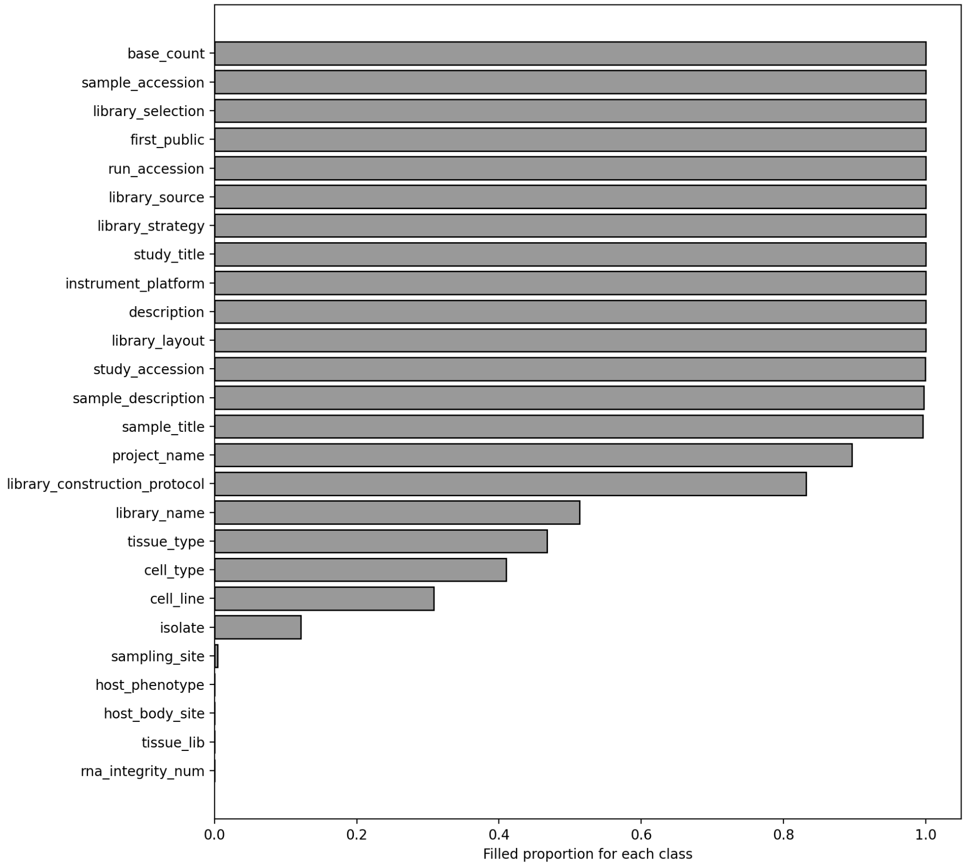


Figure 9 – **Proportion of missing SRA fields used by Metappuccino as input.** *computed over 825 476 SRA samples filtered by :* `taxon = tax_eq(9606)`, `strat = library_strategy="rna-seq"`, `dates = first_public` $\in$ `[2012-01-01, 2024-12-31]`, `plat = instrument_platform="illumina"`, `counts = read_count` $\geq$ `10 000 000`.

### 2. MetappuccinoLLModel : In-Depth Methods

***Training Idea***

`Mistral-7B-Instruct-v0.3` is the backbone model used. One *LoRA* adapter (17) is trained per class and plugged onto this backbone when needed : the class key in the prompt (e.g. `"organ"` or `"library_selection"`) deterministically selects the corresponding adapter, which is then swapped by enabling its LoRA weights on the frozen backbone (adapters are cached and activated/deactivated without reloading the base model). The prompt from Table 2 is reused. This keeps deployment fast, modular, and memory-efficient.

***Training procedure and final adapters selection***

At each *step*, the model receives the prompt plus the beginning of the JSON answer (e.g. `{"organ":"`) and is trained to predict the next token(s), i.e. the value. Reference tokens (e.g. `lung`) serve only as supervision targets (Supplementary Figure 10.1-2.). For the received prompts at each step, the model attempts to predict the answer using two components : the LM head and the classification head. For open-vocabulary classes, only the LM head, native to the Mistral's architecture, is used. It simply predicts the next tokens until a complete answer is produced. This answer is then compared to the ground truth, and a cross-entropy loss is computed on the answer. The loss is a numeric score that tells how far the model's outputs are from the correct answers. Lower loss means the model's predictions match the targets better. And a cross-entropy loss is the negative log-likelihood of the true label under the model's predicted probabilities, so it is small when the model gives high probability to the correct class and large when it does not. Then, for discrete classes a classification head was added only for the training phase to help the LM Head. It makes it possible to compute the probabilities for each value that the class can output and therefore has a certain weight in the loss depending

on the stage of training, to push it to really choose among the possible classes (Supplementary Figure 10.3-4.). Finally, depending on the outputs, the weights of the LoRA matrices are changed, and it is these weights that form the adapters the user downloads to run MetappuccinoLLModel, in addition to the original Mistral weights (Supplementary Figure 10.5.).
Validation reuses the same prompt with greedy decoding (the model selects the token with the highest probability *argmax*) and its performances are computed by soft or normal accuracy as described in the section *Performance Evaluation*. Early stopping and checkpoint selection saves the model that achieved the highest accuracy on the validation sets for each class. Hyperparameters and settings are documented in the Hugging Face configs of the released adapters (https://huggingface.co/chumphati/MetappuccinoLLModel).

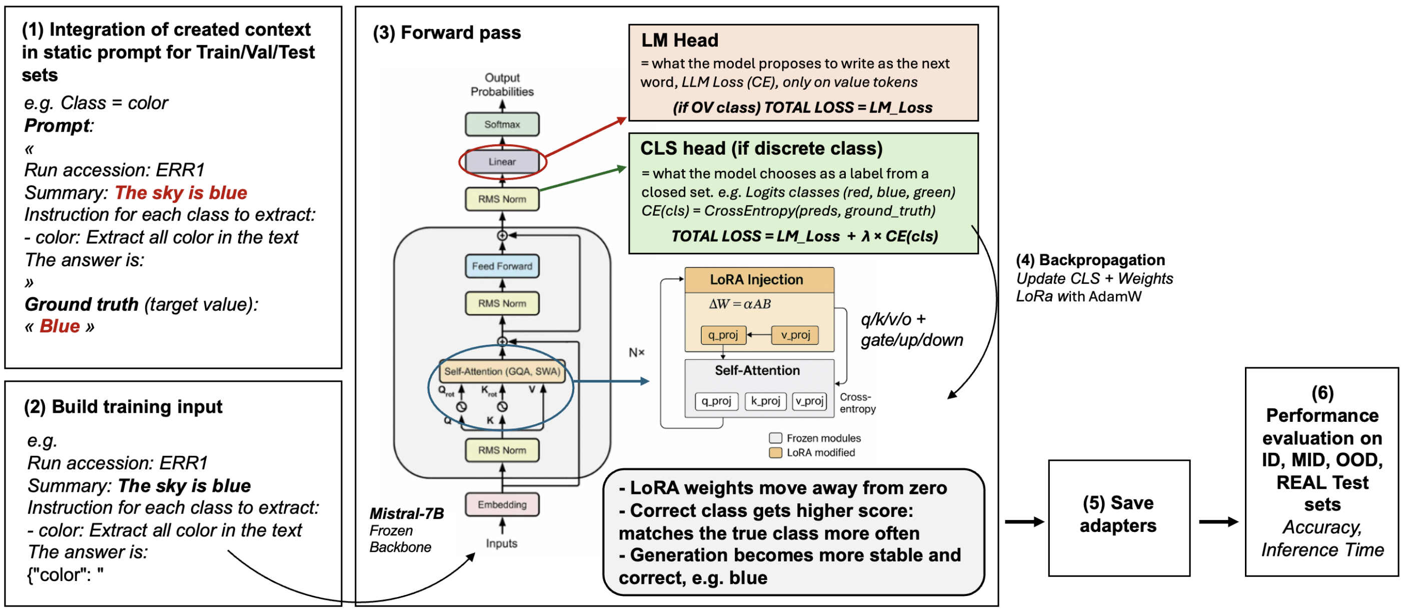


FIGURE 10 – **MetappuccinoLLModel Training Workflow.** (1–2) Inject the data into the prompt and prepare the model input. (3) Run a forward pass with the untuned Mistral-7B (frozen weights) : an LM head generates the next token, and an added classification head for discrete classes computes probabilities over a finite set of values. Ground truth is used to determine whether a prediction is good or not. (4) Backpropagation updates only the LoRA parameters, leaving the base model weights unchanged. (5) Save the updated LoRA matrices during training. (6) At inference, the original model is used and the saved adapters are activated whenever their associated class must be predicted, to assist the base model.

#### *Data Generation*

A key part of training the model and objectively evaluating its performance is to obtain a clean, unbiased dataset.

***Extracting diverse data for model training*** Since MetappuccinoLLModel's task involves extracting information that is rarely well-structured, obtaining sufficiently large, clean, and varied labeled datasets is difficult. Sentences with explicitly annotated fields tend to follow a repetitive and simplistic pattern (e.g. "class name : value"), which leads models to memorize these patterns rather than learning how to infer information from context. To overcome this limitation, synthetic data was generated. Real SRA metadata were used as the foundation, and the original values of the classes (unknown because they were not annotated) were removed while new ones were inserted. To do so, metadata text blocks were stripped of any words or expressions that could correspond to the 15 classes, resulting in contextual templates resembling authentic metadata but devoid of explicit answers. Each removed element was replaced by a placeholder indicating the class name (e.g. "CELL_LINE"), allowing controlled substitution of class-specific values (Supplementary Figure 11). This approach ensured balanced representation across all classes and produced synthetic data that accurately mirrored the linguistic structure of the real SRA metadata.

***Template generation*** Two complementary strategies were applied to construct these templates : (I) a manually curated approach (around one hundred templates, divided in 3 sets to not have the same templates for the following train, validation and tests subdivisions) available in the data section of the GitHub repository) (Supplementary Figure 11.1.) and (II) an automated approach combining rule-based methods with large language model reformulation (using Qwen to avoid bias toward Mistral's phrasing) (Supplementary Figure 11.2.). This automated procedure enables large-scale generation and reduces structural redundancy across datasets that mechanically happens with limited, manually curated data. However, this method occasionally fails to remove all ambiguous mentions (for example, implicitly suggesting bulk sequencing could be hard to remove consistently).
To generate those templates, (I) 800 000 human RNA-seq SRA metadata run accessions were collected and then deduplicated to

retain one run per study (38236 samples). Then, (II) the remaining samples were downloaded and summarized with Metappuccino preprocessing steps. (III) Those summaries were then embedded using Mistral, so each row is a normalized vector for one run. (IV) For training, we need a representative sampling of the whole pool, to build train and validation sets. We chose to generate clusters to approximate modes/themes in the population. To do that, we did a PCA (n components is 64) to denoise and make distances more meaningful. Then clusters were formed using MiniBatchKMeans on the PCA projection, using Euclidean distance. (V) From these clusters, 3000 training samples were chosen with cluster-proportional quotas method, taking mostly near-centroid points (the core) plus a smaller fraction of peripheral points to cover diversity (Supplementary Figure 11). 500 validation and 1000 tests samples were drawn from the remainder (after removing train points), matching the cluster distribution of the remaining pool, but biased slightly toward the periphery compared to train (Supplementary Figure 11). This allows to capture structurally a majority of the diversity of the known human RNA-seq metadata contexts. These 3500 samples are then automatically processed as explained previously and transformed into templates. Combined with about one hundred manually curated templates, these 3600 templates formed a base to generate good training datasets for each class.

***Value integration and dataset balancing*** To fill the templates, biologically coherent dictionaries (available on Github) were manually constructed to ensure consistency among classes (e.g. associating "lung" with a liver disease). Values were integrated to achieve balanced representations across all classes (Supplementary Figure 16, 11). Discrete classes shared identical value sets across all splits, while open-vocabulary classes contained at least twenty distinct values per class and per dataset, each associated with a minimum of five synonyms to introduce linguistic diversity. When no placeholder existed for a class, corresponding information was inserted randomly into the context. The same value could appear multiple times depending on the number of detected placeholders. For open-vocabulary classes, no value overlap existed between training, validation, and test sets, ensuring that models learned to infer meaning rather than memorize specific values.

***Per-class Train-Set & Validation-Set*** Different metadata classes responded differently to input sources. Some benefited more from automatically generated templates, whereas others performed better with manually curated data. Consequently, train and validation compositions varied by class (Supplementary Figure 14, 12.1.). `library_selection`, `biopsy_site`, `organ`, `disease`, and `sex` were trained on 2990 synthetic examples built from automatically generated templates, with 499 validation examples constructed in the same way but using distinct values. The class `biopsy_type` required additional rebalancing (blood, primary, metastasis) and therefore used 1000 training and 300 validation examples with different values but was still built with the automatic templates. In contrast, `sequencing_source`, `cell_line`, `cell_type`, `age`, `treatment`, `treatment_time`, `response`, `is_cancer`, and `ethnicity` achieved better results when trained on mixed datasets combining synthetic and real SRA contexts. Depending on data availability, 20–50% of their training data consisted of authentic SRA metadata with reliable annotations. However, 100% real data were avoided to prevent overfitting on trivially annotated patterns (e.g. "class name : value"). The remaining part of training sets and the validation sets for these classes were built from manually curated templates (duplicated to get at least 1000 training data) (Supplementary Figure 12.1.). Supplementary Figure 16 reports, for each class, the number of values used and their distribution across the train, validation, and test sets.

***Per-class synthetic Test-Sets*** To provide robust evaluation, 3 test splits were created for each class : in-distribution (ID), out-of-distribution (OOD), and middle-distribution (MID) (Supplementary Figure 12.2.). ID samples had contextual embeddings close to their associated training sets and thus were expected to yield the best performance. OOD samples were chosen for maximal distance in embedding space and represented structurally distinct contexts, where performance was expected to decline. MID samples occupied intermediate positions, providing insight into model behavior across gradual distribution shifts. Each split contained 400, 400, and 1,200 samples respectively (Supplementary Figure 14, 12.2.). For every class, two raw test sets were generated by filling the test automatic (1000) and manual (1000) templates with new equilibrated classes values (2000 total) Figure 12.2.). 30% of class values were added in a clear way (the answer pasted directly into the text), and 70% were added in a masked way by mixing in synonyms or noise so the information is harder to find. Those 2000 synthetic samples where then separated into the 3 test subsets (ID, MID, OOD) according to their embedding similarity to the training data using Mistral embeddings by separability analysis (AUC-based classification). The AUC is computed from the embeddings by (I) calculating the training centroid (the L2-normalized mean vector of the L2-normalized training embeddings), (II) measuring the cosine similarity between this centroid and the embeddings of each dataset to be compared. The higher the cosine score, the closer the dataset to compare is to the training distribution, and conversely. This is why AUC(test ID) $>$ AUC(test MID) $>$ AUC(test OOD) is expected, which is indeed observed (Supplementary Figure 14). The coexistence of manual and automatic templates naturally facilitated structural diversity : when a model was trained on automatically generated templates, manually constructed ones served as ideal OOD tests, and conversely.

***Real Test-Set*** In addition to the previously described synthetic test datasets, a real dataset was created. This set comprised 400 randomly selected SRA contexts that were disjoint from training at the run-accession level, plus 100 additional contexts from a melanoma case study that were also randomly selected. Because real SRA records do not provide complete reference labels for the 15 classes, accuracy cannot be computed. Ground truth was therefore established by manual annotation of all 500 samples using only the inputs available to Metappuccino, and no additionnal Cellosaurus dictionnaries (Figure 2). For the 100 melanoma samples, a second reference was produced by complementing the previous labels with Cellosaurus-based information consistent with Metappuccino's pre and postprocessing logic, which enabled performance evaluation of both the LLM inference and the end-to-end system (*Study case with Metappuccino : 100 melanoma samples* section).

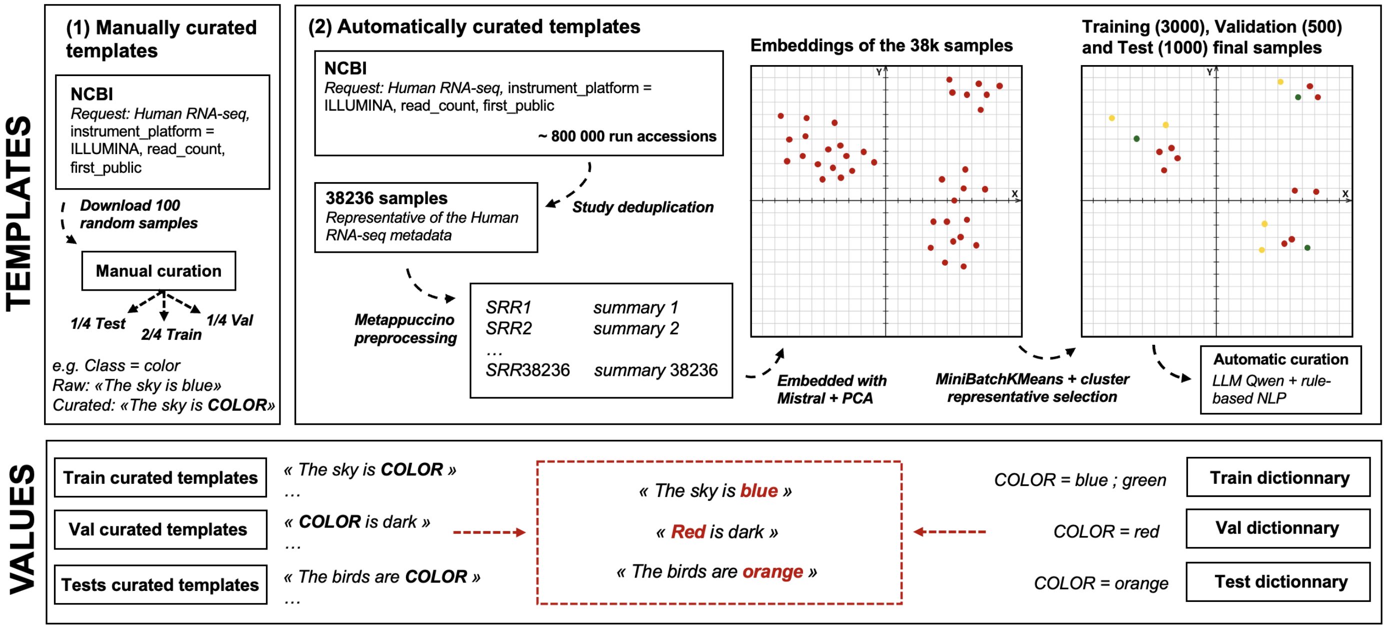


FIGURE 11 – **Methods to create the training and evaluation datasets.** Synthetic data is created from real SRA data that is cleaned either manually (1) or through automated processes (2). Automated data cleaning can introduce errors but allows for much greater diversity in the data and, above all, data that is representative of our population.

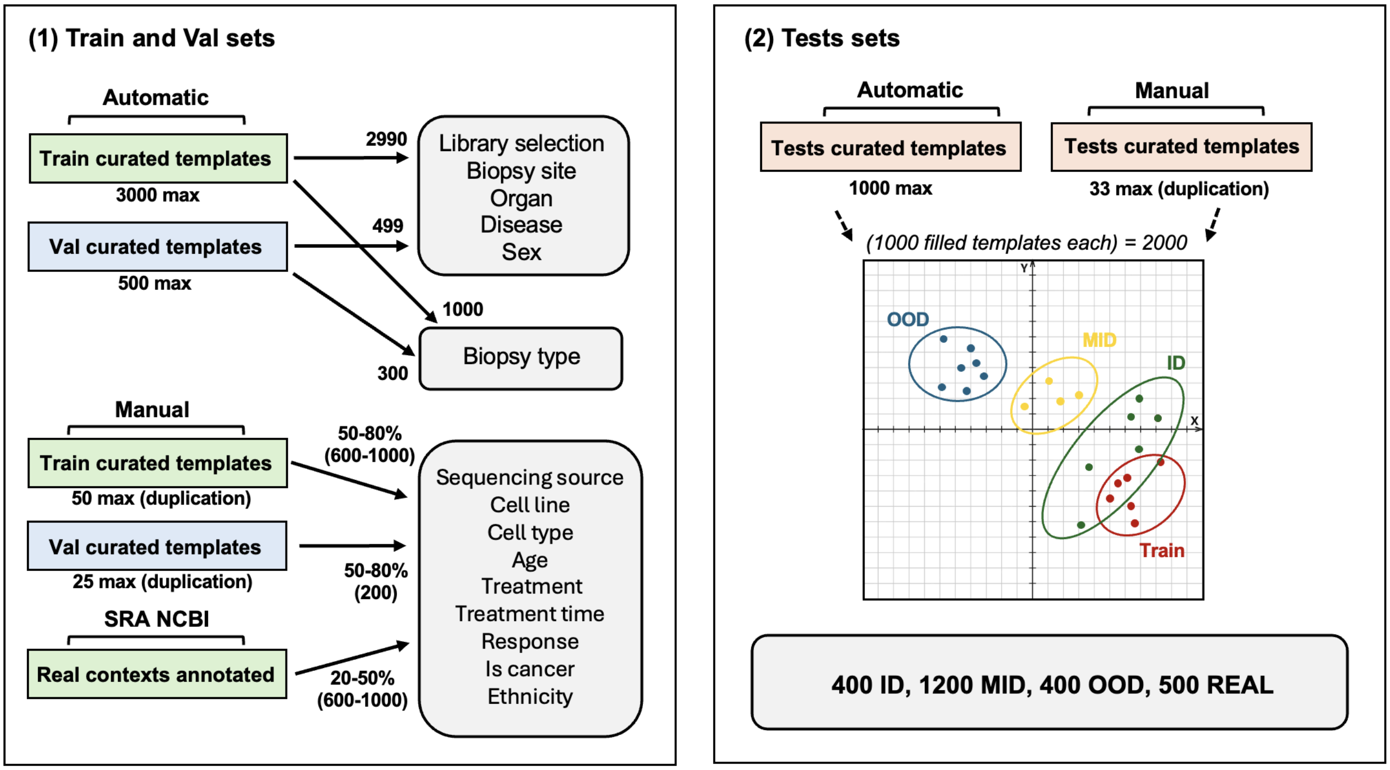


FIGURE 12 – **Dataset split methods.** (1) Each Metappuccino class is trained on distinct datasets composed of templates generated by different methods, selected based on how well they capture the target signal. This figure reports, for each class, the distribution of template types across the training and validation splits. Numbers on the arrows indicate the count or proportion of each template type, and the number beneath each template block gives the total number of templates of that type produced by the methods in Figure 11. (2) Test-set split. From each template type, 1000 samples were generated, so 2000 test samples in total. For each class, each sample's distance was computed to its own training set : the 400 nearest are labeled ID, the 1200 in the middle are MID, and the 400 farthest are OOD.

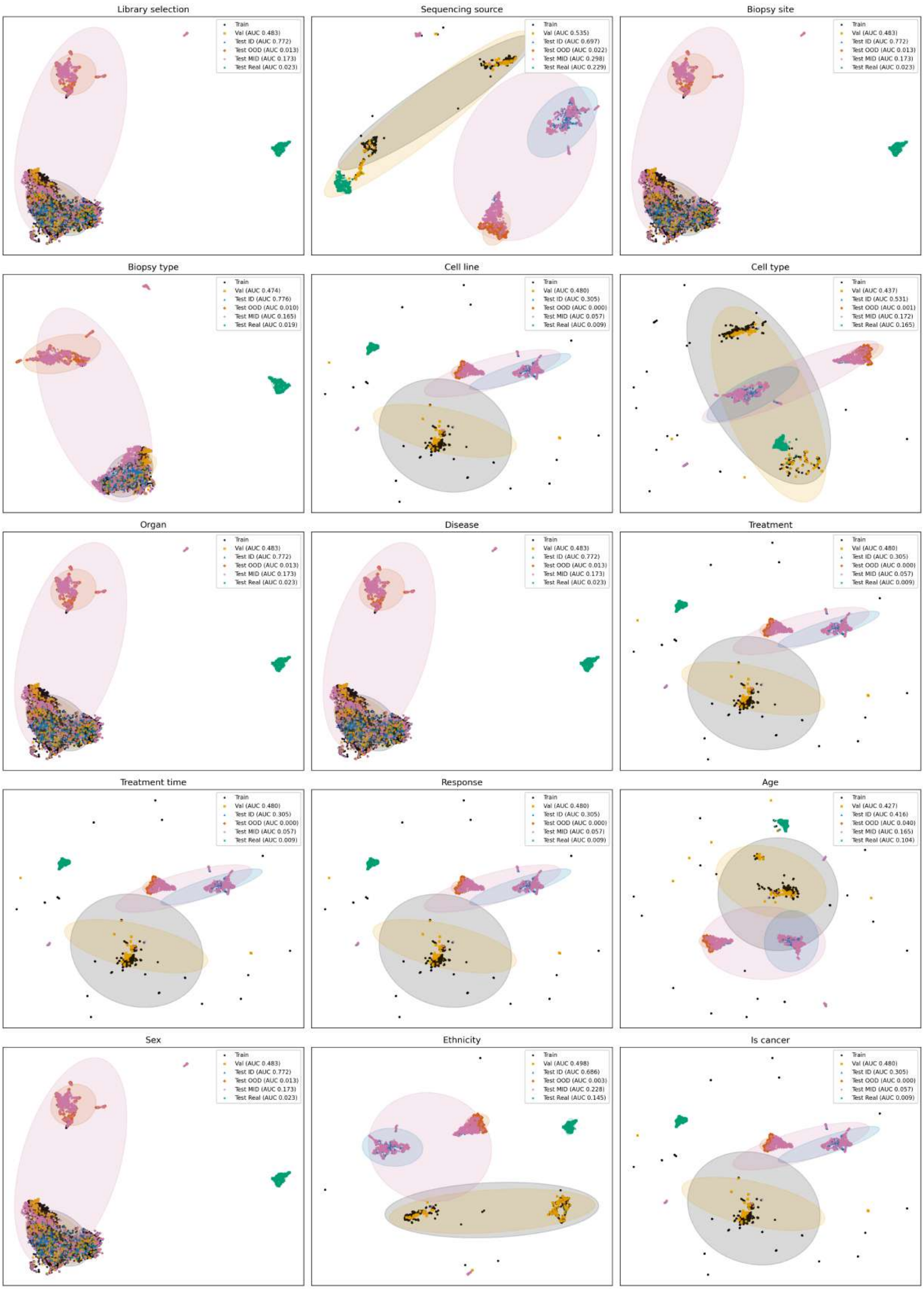


FIGURE 14 – **Embedding distributions across all metadata classes.** Two-dimensional UMAP projections of the L2-normalized Mistral embeddings for training (black), validation (orange), and test splits (ID = blue, MID = pink, OOD = red) plus real SRA contexts (green). Ellipses represent confidence regions. The AUC values indicate cosine-based separability between the training centroid and each split : higher AUC reflects stronger similarity. MID and ID sets show moderate overlap, while OOD and real contexts form distinct clusters, confirming controlled distribution shifts and minimal overlap across splits.

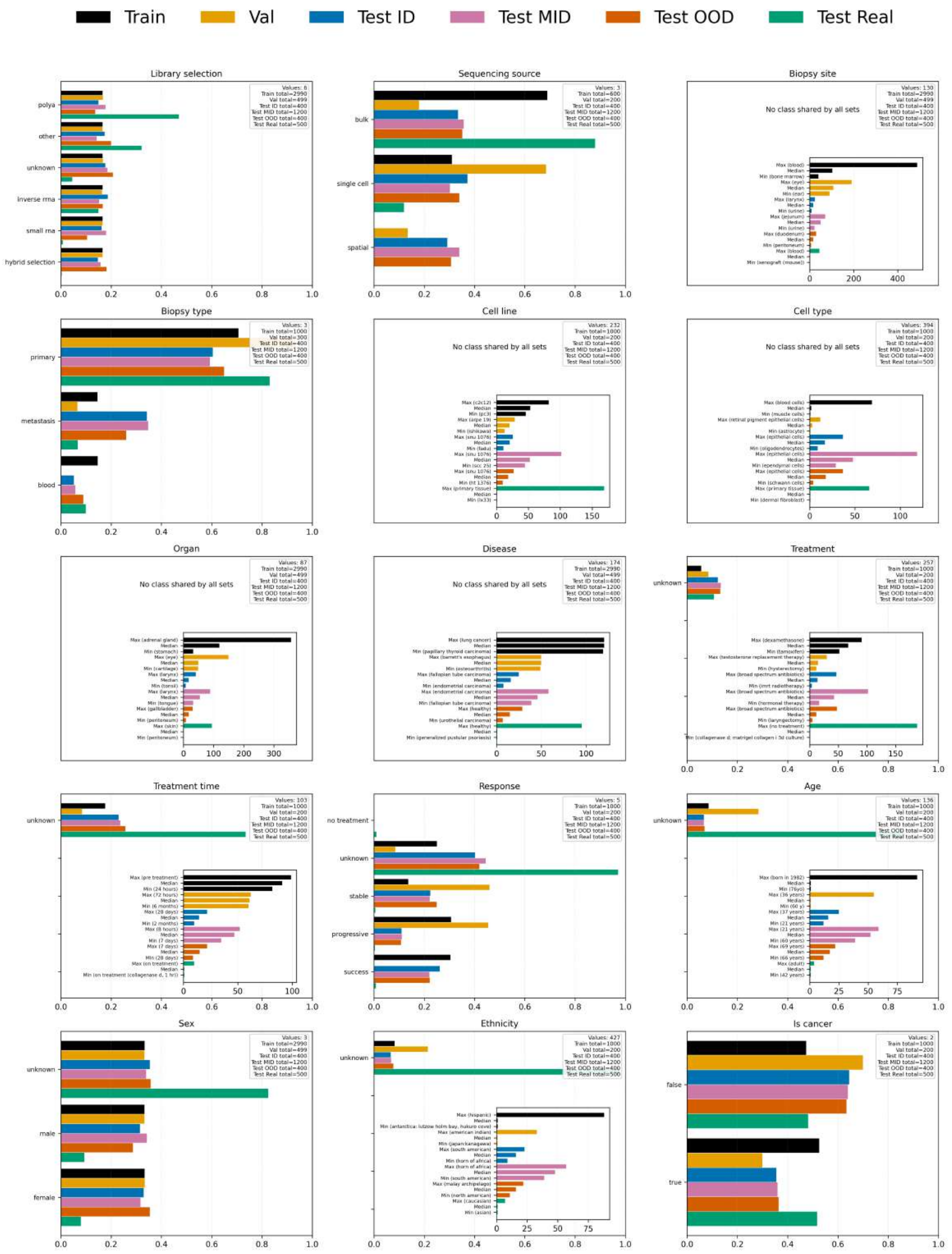


Figure 16 – **Class distributions across all metadata classes.** Distribution of class values per split (Train, Validation, Test ID, Test MID, Test OOD, and Test Real) for each metadata class. Synthetic datasets were filled using biologically coherent dictionaries to ensure internal consistency and were balanced as evenly as possible across splits through controlled template generation. Open-vocabulary classes include around 20 distinct synthetic values per dataset. Classes displaying a larger number of values correspond to those partially trained on real SRA metadata, where authentic annotations were integrated into the training and validation sets. Thoses values are listed in a JSON file available in the data section of the Github project. When perfect balance was not achievable, training algorithms were adjusted to handle the imbalance classes.

### 2. Metappuccino

***Extracted fields from the NCBI API during pipeline download*** The downloaded BioSample XML file contains sample-level record with attributes under the corresponding tag, with keys taken from `attribute_name` or `harmonized_name`) to capture details often missing at run level. In parallel, a call to the SRA API provides the following fields : `study_accession`, `first_public`, `study_title`, `project_name`, `run_accession`, `sample_accession`, `sample_title`, `sample_description`, `library_name`, `library_selection`, `library_source`, `library_strategy`, `library_construction_protocol`, `library_layout`, `rna_integrity_num`, `instrument_platform`, `rt_prep_protocol`, `cell_line`, `cell_type`, `tissue_lib`, `tissue_type`, `host_phenotype`, `isolate`, `age`, `host_body_site`, `sampling_site`, `base_count`, `description`.


```json
{
  "SRR28878223": {
    "library_selection": "other",
    "sequencing_source": "bulk",
    "biopsy_site": "lymph node",
    "biopsy_type": "primary",
    "cell_line": "not applicable",
    "cell_type": "lymph node",
    "organ": "lymph node",
    "disease": "melanoma",
    "treatment": "vidutolimab",
    "treatment_time": "pre",
    "response": "success",
    "age": "75",
    "sex": "male",
    "ethnicity": "caucausian",
    "is_cancer": "true"
  },
  "nll": {
    "library_selection": 0.0005165196489542723,
    "sequencing_source": 0.033508021384477615,
    "biopsy_site": 0.0008475284627991186,
    "biopsy_type": 6.726217091083527,
    "cell_line": 0.0035979782696813345,
    "cell_type": 0.10822681952413404,
    "organ": 6.517138845651971e-05,
    "disease": 1.1801512298366864e-05,
    "treatment": 0.5367327570915222,
    "treatment_time": 0.04310770332813263,
    "response": 0.013448241166770458,
    "age": 0.14632034579699393,
    "sex": 7.152531907195225e-06,
    "ethnicity": 0.17829522117972374,
    "is_cancer": 0.23145367205142975
  },
  "ppl": {
    "library_selection": 1.0005166530681984,
    "sequencing_source": 1.0340757384120878,
    "biopsy_site": 1.000847887716532,
    "biopsy_type": 833.9863965074119,
    "cell_line": 1.00360445876339,
    "cell_type": 1.1143004618253665,
    "organ": 1.0000651735121575,
    "disease": 1.0000118015819366,
    "treatment": 1.7104094000470738,
    "treatment_time": 1.0440503364913125,
    "response": 1.0135390754925788,
    "age": 1.1575669503688402,
    "sex": 1.0000071525574865,
    "ethnicity": 1.1951781110486497,
    "is_cancer": 1.2604309320455762
  }
}
```


FIGURE 17 – **Example of raw JSON output after LLM inference : SRR28878223.** The file has three top-level keys : the run name, nll and ppl. nll and ppl reflect model confidence per predicted class (lower is better). It reports results for a single sample at inference time ; these values are later merged by Metappuccino with normalization and pre/postprocessing into the final output.
